# What if we perceive SARS-CoV-2 genomes as documents? Topic modelling using Latent Dirichlet Allocation to identify mutation signatures and classify SARS-CoV-2 genomes

**DOI:** 10.1101/2020.08.20.258772

**Authors:** Sunil Nagpal, Divyanshu Srivastava, Sharmila S. Mande

## Abstract

Topic modeling is frequently employed for discovering structures (or patterns) in a corpus of documents. Its utility in text-mining and document retrieval tasks in various fields of scientific research is rather well known. An unsupervised machine learning approach, Latent Dirichlet Allocation (LDA) has particularly been utilized for identifying latent (or hidden) topics in document collections and for deciphering the words that define one or more topics using a generative statistical model. Here we describe how SARS-CoV-2 genomic mutation profiles can be structured into a ‘Bag of Words’ to enable identification of signatures (topics) and their probabilistic distribution across various genomes using LDA. Topic models were generated using ~47000 novel corona virus genomes (considered as documents), leading to identification of 16 amino acid mutation signatures and 18 nucleotide mutation signatures (equivalent to topics) in the corpus of chosen genomes through coherence optimization. The document assumption for genomes also helped in identification of contextual nucleotide mutation signatures in the form of conventional N-grams (e.g. bi-grams and tri-grams). We validated the signatures obtained using LDA driven method against the previously reported recurrent mutations and phylogenetic clades for genomes. Additionally, we report the geographical distribution of the identified mutation signatures in SARS-CoV-2 genomes on the global map. Use of the non-phylogenetic albeit classical approaches like topic modeling and other data centric pattern mining algorithms is therefore proposed for supplementing the efforts towards understanding the genomic diversity of the evolving SARS-CoV-2 genomes (and other pathogens/microbes).

## 1. INTRODUCTION

A document is a thematic body of text containing a semantic structure of words. The theme of a document, also called as the primary topic, is constituted by a specific proportion of various words. Considering existence of a finite vocabulary, different proportions of words (and their semantic similarity) in each document would drive the theme(s) or topic(s) of various documents. Therefore, while words are apparent constituents of a document, topics are latent (or hidden). Topic modeling, a statistical method, employs these characteristics of documents to discover hidden structures (or latent topics)^1^. Its utility in text-mining and document retrieval/classification tasks in various fields of scientific research is rather well known^2–4^. In fact, Latent Dirichlet Allocation (LDA), an unsupervised machine learning approach, is particularly known for identifying latent topics in large document collections and deciphering the words that define the inferred topics using a generative statistical model. LDA assumes that a document is generated by a distribution of all possible hidden topics, while a topic is generated by the distribution of all possible apparent words. This multiplicity of topic affiliation for documents and words is accommodated through assumption of Dirichlet priors which can be optimized to get ideal distribution of coherent topics in a document^1^. The approach can also be made akin to Markov-chains for probing the temporal evolution of a large number of documents and document topics^4^.

A large number of SARS-CoV-2 genome sequences are being deposited to public repositories like GISAID^6^ through an unprecedented spirit of scientific collaboration across the world. The high volume of raw data is expected to balloon further by the end of this pandemic. Each new sequenced genome is a mutant/variant (with few exceptions) of original reference genome i.e. Wuhan/WIV04/2019 (*EPI_ISL_402124*). In other words, certain mutations at nucleotide and amino acid levels can be expected to be observed in the submitted genomes. Understanding the evolution and diversity of these variants has been a subject of interest to a wide spectrum of researchers. Various reports aimed at identification of clades or classification system(s) for these genomes have in fact been outcomes of the afore-mentioned problem statement^7^.

Although conventional topic modeling has been utilized to understand Covid-19 from literature data^5^, can it be applied to obtain additional insights from sequenced SARS-CoV-2 genomes and to rather classify them? In other words, can we perceive each genome of SARS-CoV-2 as a document containing words in the form of characteristic mutations (Figure 1)?

**Figure 1:**
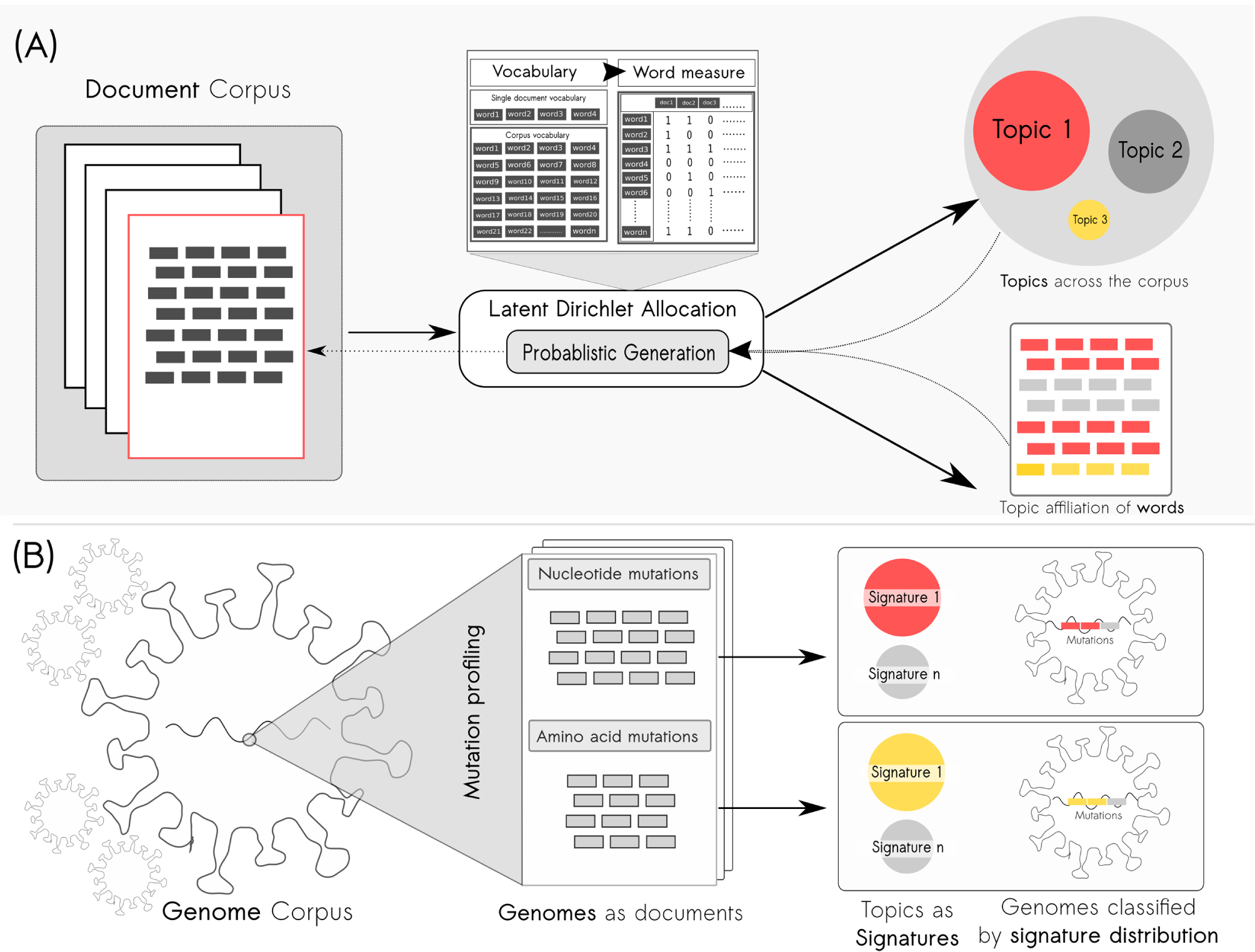
Perceiving SARS-CoV-2 genomes as documents. **Panel A:** Classical approach towards topic modeling on large document corpus using the generative process of Latent Dirichlet Allocation (LDA). **Panel B:** Each SARS-CoV-2 genome with its mutation profile is treated as a document containing words in the form of their mutations with a potential to infer latent mutation signatures (topics)

Consequently, a genome would essentially become a bag of mutations (like bag of words in a document). Such an assumption can potentially enable classification of the entire genome corpus by identifying mutation signatures (equivalent to topics in document) through topic modeling. Moreover, given the inherent temporal nature of genome collections, dynamic topic modeling (e.g. temporal LDA or Hidden Markov Model driven LDA) may rather provide a way to probe the evolution of the genome variants in terms of identified mutation signatures^4,8^. A parallel between classical LDA on a large document corpus and a genome corpus is illustrated in plate notation below.

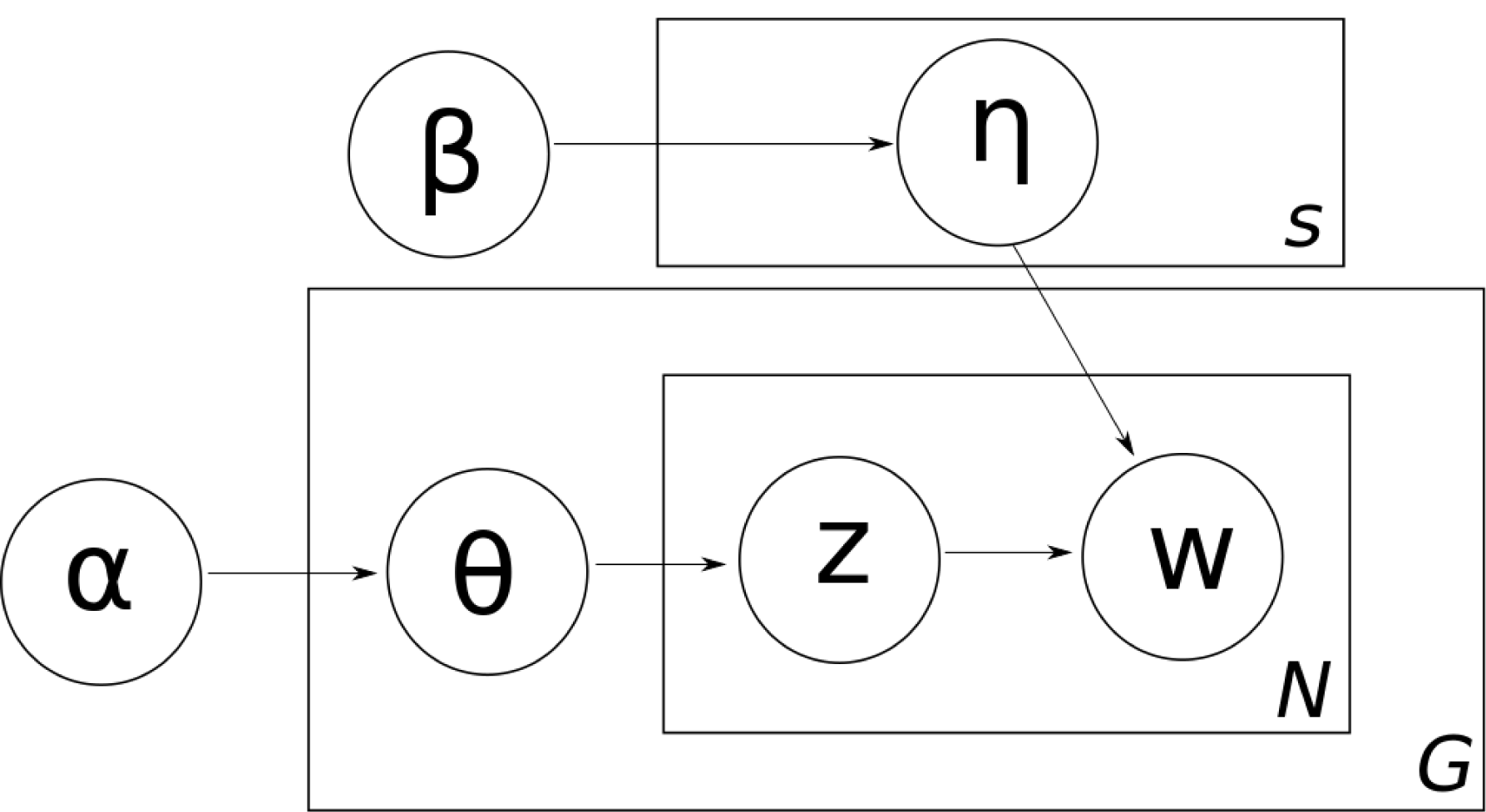

where,

**α** is a parameter governing the distribution structure of signatures (nucleotide and amino acid mutations) across all genomes (similar to topics across all documents)

**θ** is a random matrix representing Dirichlet distribution of various signatures in the genomes (similar to topics in documents), such that θ(i,j) indicate the probability of the *i* th genome (document) to contain mutations (words) pertaining to the *j* th signature (topic)

**β** is a parameter governing the distribution structure of mutations across all signatures (similar to words across all topics)

**η** is a random matrix representing Dirichlet distribution of various mutations in signatures (similar to words in topics), such that η (i,j) indicate the probability of the *i* th signature (topic) to contain the *j* th mutation (word)

**z** is an identity of signature (topic) of all mutations (words) in all genomes (documents)

**w** refers to identity of all mutations (words) in all genomes (documents)

**G** refers to all genomes (documents)

**N** refers to all mutations (words)

**S** refers to all signatures (topics)

which may be interpreted as following:

1. For each signature (topic) s, draw η_s_∼Dirichlet(β)
2. For each genome (document) g, first draw θ_g_∼Dirichlet(α), then for each *n* th mutation (word) of the genome (document) g, draw z_gn_∼Multinomial(1,θ_g_) followed by w_gn_∼Multinomial(1,η_zgn_)

To substantiate the conjecture, a bag of mutations data structure for ~47000 SARS-CoV-2 genomes submitted to GISAID was created. Classical LDA was employed to generate topic models leading to identification of 16 amino acid mutation signatures and 18 nucleotide mutation signatures (equivalent to topics) in the corpus of chosen genomes through rigorous hyper-parameter tuning for coherence optimization (Figure 2). Interestingly, most of the high weight inferred signatures had a good overlap with the previously identified clades specific to various geographical regions (**refer** Table 1). For example, the signature-11, constituted predominantly by amino acid mutations N-P13L/ORF9b-P10S, ORF1a-L3606F, ORF1b-A88V, ORF1a-T2016K, was observed to dominate in India and other Asian regions^9^. Biology agnostic, data structure driven approaches for SARS-CoV-2 genome sequences may therefore have some merit in not only handling the large amount of genomic data, but also for identifying mutation signatures (and hence classifying genomes) that might be of interest to clinicians/ biologists^10^. Their cross validation against phylogenetic estimations can help fine tune the performance of these machine learning algorithms, thereby adding confidence to the use of unconventional methods for probing genomic diversity^7^.

**Figure2:**
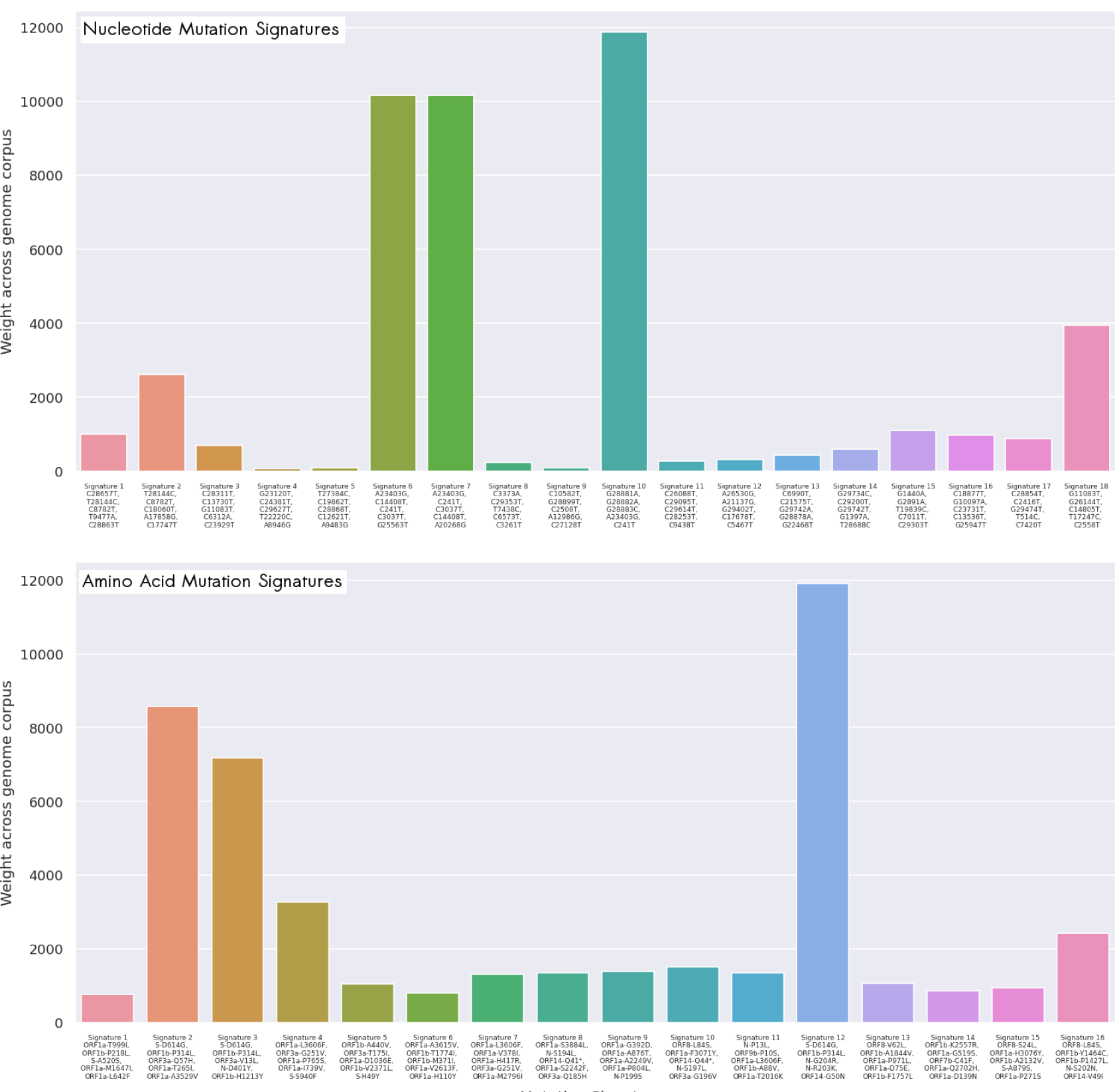
Mutation signatures in SARS-CoV-2 genomes. Nucleotide and Amino acid mutation signatures identified through classical LDA and their weights across genome corpus

**Table 1.** Putative amino acid mutation signatures, their weight across genome corpus and closest literature citing the said mutation(s). Probability of co-occurrence of these mutations in a signature was found to be low in low scoring signatures.

|  | **Mutation signature (LDA derived)** | **Score**<br>**(cumulative**<br>**weight across**<br>**genomes)** | **Reference** |
| --- | --- | --- | --- |
| **1.** | ORF1a-T999I, ORF1b-P218L, S-A520S, ORF1a-M1647I, ORF1a-L642F | 762 | No evidence |
| **2.** | **S-D614G, ORF1b-P314L, ORF3a-Q57H, ORF1a-T265I, ORF1a-A3529V** | **8581** | [16] |
| **3.** | **S-D614G, ORF1b-P314L, ORF3a-V13L, N-D401Y, ORF1b-H1213Y** | **7171** | [17] |
| **4.** | **ORF1a-L3606F, ORF3a-G251V, ORF1a-P765S, ORF1a-I739V, S-S940F** | **3280** | [18] |
| **5.** | ORF1b-A440V, ORF3a-T175I, ORF1a-D1036E, ORF1b-V2371L, S-H49Y | 1051 | [19] |
| **6.** | ORF1a-A3615V, ORF1b-T1774I, ORF1b-M371I, ORF1a-V2613F, ORF1a-H110Y | 801 | [20] |
| **7.** | ORF1a-L3606F, ORF1a-V378I, ORF1a-H417R, ORF3a-G251V, ORF1a-M2796I | 1297 | [21] |
| **8.** | ORF1a-S3884L, N-S194L, ORF14-Q41*, ORF1a-S2242F, ORF3a-Q185H | 1350 | [19] |
| **9.** | ORF1a-G392D, ORF1a-A876T, ORF1a-A2249V, ORF1a-P804L, N-P199S | 1389 | [16] |
| 10. | ORF8-L84S, ORF1a-F3071Y, ORF14-Q44*, N-S197L, ORF3a-G196V | 1507 | [22] |
| 11. | N-P13L, ORF9b-P10S, ORF1a-L3606F, ORF1b-A88V, ORF1a-T2016K | 1351 | [9] |
| 12. | **S-D614G, ORF1b-P314L, N-G204R, N-R203K, ORF14-G50N** | **11912** | [23] |
| 13. | ORF8-V62L, ORF1b-A1844V, ORF1a-P971L, ORF1a-D75E, ORF1b-F1757L | 1064 | [24] |
| 14. | ORF1b-K2557R, ORF1a-G519S, ORF7b-C41F, ORF1a-Q2702H, ORF1a-D139N | 854 | No evidence |
| 15. | ORF8-S24L, ORF1a-H3076Y, ORF1b-A2132V, S-A879S, ORF1a-P271S | 950 | [24] |
| 16. | ORF8-L84S, ORF1b-Y1464C, ORF1b-P1427L, N-S202N, ORF14-V49I | 2420 | [23] |

## 2. METHODS

### 2.1. Mutation profiles

Approximately 47000 SARS-CoV-2 sequences, obtained from Global Initiative on Sharing Avian Influenza Data (GISAID) between Jan-July 2020, were used. NextStrain’s Augur pipeline was employed with default parameters to align the sequences against the reference Wuhan/WIV04/2019 (*EPI_ISL_402124*)^11^. Individual proteins of SARS-CoV-2 were extracted post alignment and translated to the amino acid sequences. Comparisons to reference amino acid and nucleotide sequences were performed to profile mutations for all viral genome sequences. A genome collection (document corpus) mapped to the identified nucleotide and amino acid substitution mutations (document vocabulary) was created. Sample mutation profile data structure have been provided in **Supplementary Table 1.** It may be noted that only those genomes were employed for topic modelling which contained at least one amino acid mutation.

### 2.2. Bag of mutations

As shown in Figure 1, bag of words representation of a document in natural language processing pertains to two aspects of the document:

i. Document vocabulary **(V)** represented by all words of the given document
ii. Token/Word measure **(W)** represented by the occurrence profile of words in the document

With an aim to develop a ‘bag of words’ model, the mutation profiles of SARS-CoV-2 genomes used in this study was compiled such that the individual genome-specific nucleotide and amino acid mutation vocabularies (set of mutations in a genome) could be easily comprehended **(Supplementary Table 1)**. Two corpus vocabularies were consequently created (one each for nucleotide and amino acid mutations). Binary document vectors were prepared for each of the genomes against the corpus vocabularies for these two types of mutations. Mutation-genome matrices so computed for the two corpora represented the global picture of ‘bag of words’ models for novel corona virus genomes. It may be noted that unlike a conventional natural language processing task, given the non-linguistic context of observed mutations, the issues pertaining to tokenization, stop words, lemmatization and stemming were not relevant here^1^.

#### 2.2.1. Bag of mutation bi-grams

Given that most of the existing clade definitions employ two or more co-occurring mutations, a bigram nucleotide mutation model was also created for the genomes. The corpus vocabulary for bigrams was created by taking into account the observed co-occurring pairs of mutations in the entire corpus of nucleotide mutation vocabulary (and not all possible pairs of mutations), such that each genome was represented by a numerically sorted list of nucleotide mutations. It is pertinent to note that numeric sorting of mutations is critical in searching for bi-grams (or n-grams) for a meaningful contextual search. It may be noted that the choice of bi-gram mutations is enabled by a probabilistic scoring procedure^12^ as follows:

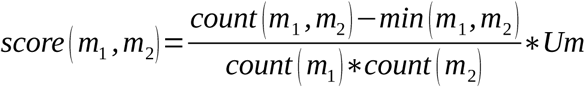

where:

**m_1_,m_2_** are mutations in a pair

**Um** is total unique mutations (i.e. size of mutation vocabulary)

**score** refers to the confidence score for the given pair

**count** refers to the total occurrence in the corpus

**min** refers to the minimum occurrence threshold for the mutation(s) in the corpus

#### 2.2.2. N-gram mutation signatures

The probabilistic derivation of bi-grams paves the way for an initial estimation of signatures of any size in the corpus using the following progressive probabilistic scoring:

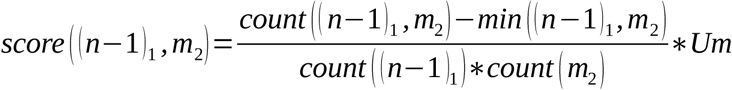

where:

**(n-1)_1_, m_2_** are mutations in a pair, such that (n-1) refers to the (n-1) sized mutation combination

**Um** is total unique mutations (i.e. size of mutation vocabulary)

**score** refers to the confidence score for the given pair

**count** refers to the total occurrence in the corpus

**min** refers to the minimum occurrence threshold for the mutation(s) in the corpus

### 2.3. Topic (mutation signatures) modeling through Latent Dirichlet Allocation and hyper-parameter tuning

Python’s Gensim library was employed to estimate topic (mutation signature) models for ~47000 SARS-CoV-2 genomes through online variational Bayes (VB) algorithm as described previously^13,14^. Quality of mutation signatures (topics) inferred by LDA was assessed using a coherence score which refers to an index of the semantic similarity between dominant mutations (words) of the mutation signature (topic). In other words, a mutation signature (topic) with high coherence is expected to have mutations (words) with high co-occurrence similarity score. A good overall mutation signature extraction is therefore expected to have a high mean coherence. The coherence measure was calculated for different numbers of mutation signature extractions between 2-30 and an optimal score for nucleotide as well as amino acid mutations were obtained. Further hyperparameter optimization was performed for a range of alpha and beta measures (between 0.001 – 0.1, step size of 0.009) and the number of topics, in order to maximize the coherence score, and optimal values for all three parameters were obtained using the grid-search alogrithm^15^.

### 2.5 Implementation

The entire implementation was executed in a 20 core Xeon 51 series 2.4GHz machine with 64GB RAM in a Python v3.7.6 kernel with Gensim v3.8.3 and Scikit-learn v0.23.1 for topic modelling using LDA.

## 3. RESULTS

### 3.1. Word clouds of the corpus-wide bag of mutations

Word clouds provide quick visual reference to the dominant words in a bag of words. As shown in Figure 3a, the bag of nucleotide mutations for all genomes indicated the dominance of A23403G, C14408T, G28881A, G28882A, G28883C, C3037T and C241T amongst the 15114 unique mutations observed in the entire corpus (359976) vocabulary (all nucleotide mutations in the genomes). Similarly, among the total 9583 unique amino acid mutations, predominant ones pertained to those in Spike (S): D614G, ORF1b: P314L, ORF14: G50N and Nucleocapsid (N): R203K, G204R (Figure 3b). This approach therefore provided a preliminary way of quickly visualizing the global mutation signatures.

**Figure 3:**
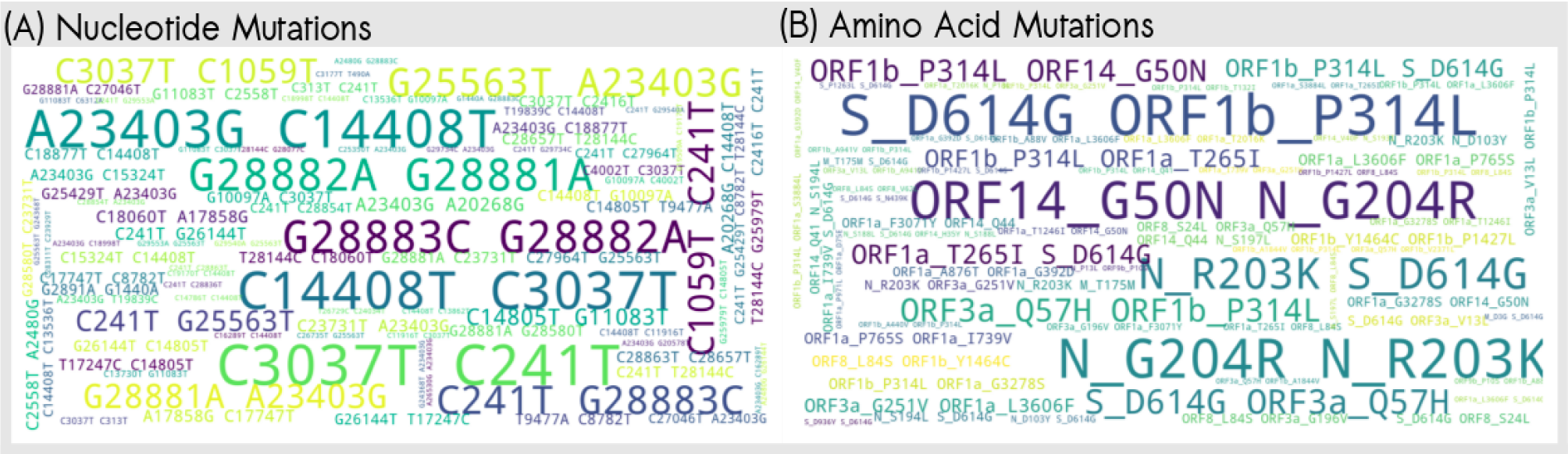
Mutation word clouds. Word clouds of the corpus-wide bag of (A) nucleotide and (B) amino acid mutations

### 3.2. Bi-grams and tri-grams

At a minimum co-occurrence count of 500 genomes and threshold of 1, 28 nucleotide mutation bigrams were identified, the most frequent (in ~13000 genomes) bi-grams being G28882A_G28883C and G28881A_G28882A, followed by A23403G_G25563T (9282 genomes). **Supplementary Table 2** provides a full list of the detected bi-grams along-with their respective scores and co-occurrence counts. Similarly, **Supplementary Table 3** provides a list of 12 tri-grams. G28881A_G28882A_G28883C was observed to be the most frequent tri-gram (12730 genomes), followed by A23403G_G28881A_G28882A (6928 genomes) and C241T_C1059T_C3037T (6556 genomes). The probabilistic approach can be extended to co-occurring contextual mutations of any size (n-gram) (described in methods section).

### 3.3. Mutation signature identification and genome classification using Latent Dirichlet Allocation

16 amino acid mutation signatures and 18 nucleotide mutation signatures were obtained at an alpha (α) value of 0.005 and beta (β) 0.067. These hyper-parameters, as described in the Methods section, were optimized through grid search. Figure 2 provides an overview of the Top 5 mutations constituting each signature and the distribution trend of the signatures across various genomes. Given that each signature has a Bayesian probabilistic estimate of occurrence in a genome, the dominant signature of each genome was looked for. This enabled the classification of each genome in terms of its dominant signature affiliation. A world map visualization of the sampling location of each genome and its signature affiliation helped in obtaining an intuition regarding the global diversity and spread of SARS-CoV-2 genomes (Figure 4).

**Figure 4:**
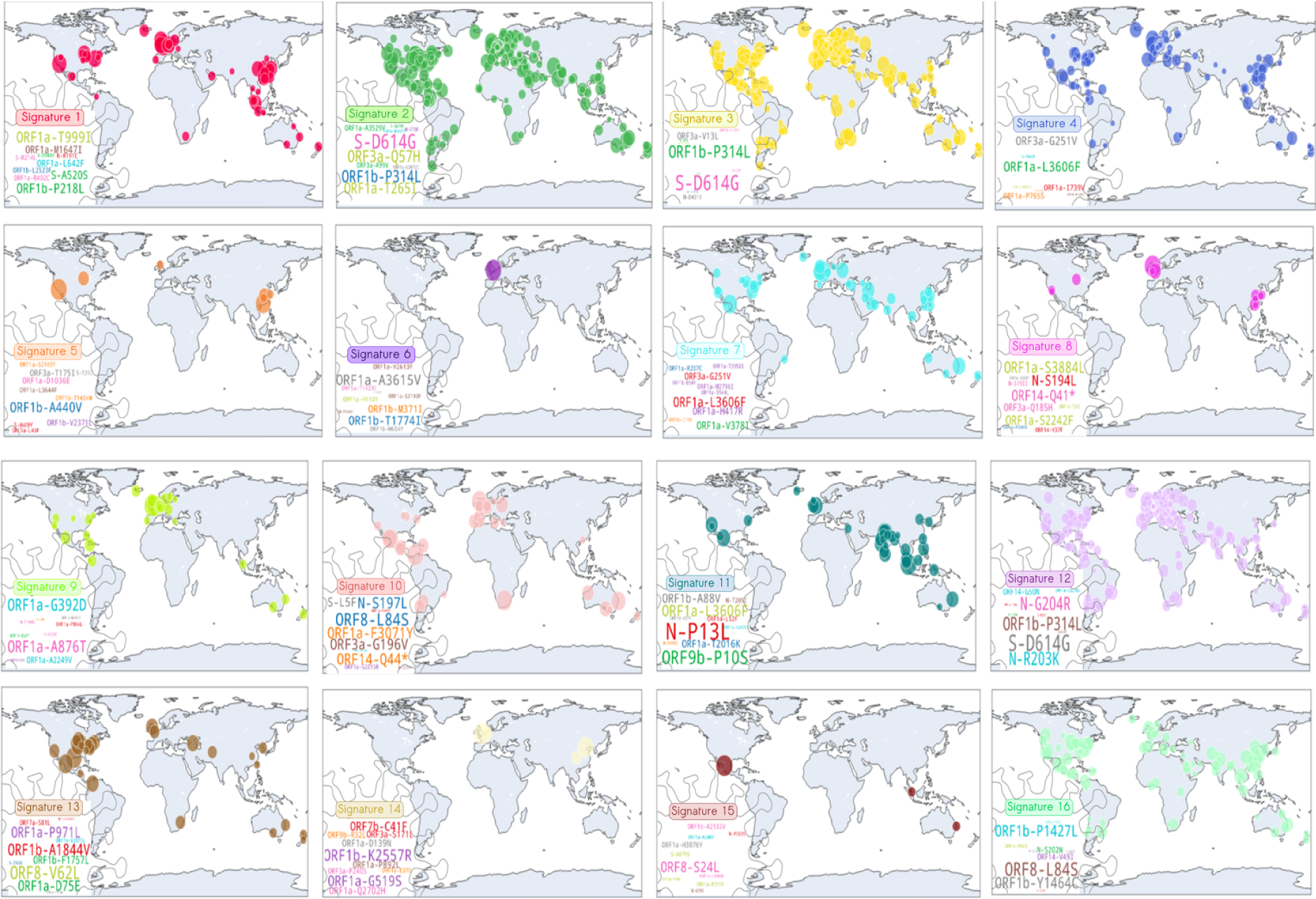
Geographical spread of putative signatures. Global map of geographical spread of putative amino acid signatures

### 3.4. Validation of mutation signatures

Validating non-phylogenetic algorithms of genome classification against phylogenetic estimations can provide an index of suitability of the data structure driven methods. As a qualitative cross-checking, the dominating mutation composition of signatures inferred using LDA was compared with the well known recurrent mutation reports and clade definitions. Table 1 provides a summary of the amino acid mutation signatures detected through LDA and corresponding close literature evidence citing a similar phylogenetically estimated genome group/clade (if any). The mutations in the signature were ordered in according to the probability of their presence in the signature. Consequently, each signature was dominated by the first mutation, as compared to the probability of occurrence of other mutations in the signature. Also, it is pertinent to note that given the probabilistic nature of inference, a high total score (weight) is more likely to indicate co-occurring mutations across large number of genomes. First five mutations, in the order of their probability of occurrence in the signatures, have been listed in Table 1. In addition, the bi-grams and tri-grams identified through probablistic approach in this study have already been supported with their score and prevalence across genomes (**Supplementary Table 2 and 3**).

## Discussion

While the mutation signatures obtained through unsupervised machine learning approaches are not phylogenetic, right choice of algorithms can enable identification of probabilistic (and hence reliable) markers to classify genomes based on observed mutations. In fact, an evolutionary trail may also be established by following a temporal approach to LDA (or other methods of topic modeling). An increase in efficiency of signature detection may further be achieved through other topic modeling methods (e.g. short text topic modeling). Importantly, insights obtained about latent signatures through machine learning approaches like LDA can also guide phylogenetic estimations. This article is intended to encourage the use of unconventional data driven approaches as an avenue that deserves attention of both data scientists and biologists alike. This, we believe, is expected to supplement the efforts in understanding the genomic diversity of the evolving SARS-CoV-2 genomes (and other pathogens).

## Acknowledgement

We gratefully acknowledge all the Authors from the Originating laboratories responsible for obtaining the specimens and the Submitting laboratories where genetic sequence data were generated and shared via the GISAID Initiative, on which this research is based. Genome sequences and meta-data should be downloaded from https://www.gisaid.org. A sample file for mutation profiles generated for this research has been provided in Supplementary Table 1 along with the original contributors of these virus genome sequences in Supplementary Table 6. Authors would also like to thank the management of Tata Consultancy Services Ltd for promoting the environment of fundamental and applied research. Authors would like to thank their colleague Nishal K. Pinna for his assistance in generating mutation profiles.

## Author contribution

SN conceived the idea and designed the study. SN and DS performed the analyses. SN, DS and SSM analysed the results. SN wrote first draft of manuscript and designed figures. SSM supervised the work and finalized manuscript. All authors reviewed and approved the submission.

