## Supplementary Table 1 for "What if we perceive SARS-CoV-2 genomes as documents? Topic modelling using Latent Dirichlet Allocation to identify mutation signatures and classify SARS-CoV-2 genomes"

**Supplementary Table 1: Sample Mutation profile data structure to enable creation of individual genome-specific nucleotide and amino acid mutation vocabularies**

| Gisaid_EPI_ISL_ID | Nucleotide_Mutations | Amino_AcidMutations |
|---|---|---|
| EPI_ISL_418241 | A23403G,C10582T,C1059T,C14408T,C18115T,C241T,C25777T,C26461T,C29353T,C3037T,G25563T | E-L73F,ORF1a-T265I,ORF1b-H1550Y,ORF1b-P314L,ORF3a-L129F,ORF3a-Q57H,S-D614G |
| EPI_ISL_418242 | A23403G,C10582T,C1059T,C14408T,C241T,C29353T,C3037T,G25563T | ORF1a-T265I,ORF1b-P314L,ORF3a-Q57H,S-D614G |
| EPI_ISL_420037 | A23403G,C10582T,C1059T,C14408T,C241T,C3037T,C5730T,G25563T | ORF1a-T1822I,ORF1a-T265I,ORF1b-P314L,ORF3a-Q57H,S-D614G |
| EPI_ISL_452463 | C14805T,C28657T,C28863T,C8782T,G25979T,G29402T,G29742T,T28144C,T9477A | N-D377Y,N-S197L,ORF14-Q44*,ORF1a-F3071Y,ORF3a-G196V,ORF8-L84S |
| EPI_ISL_413485 | A5094G,C16049T,C2189T,C8782T,G17122A,T28144C,T3086C | ORF1a-F941L,ORF1a-H1610R,ORF1a-L642F,ORF1b-A1219T,ORF1b-T861I,ORF8-L84S |
| EPI_ISL_420600 | A23403G,C14408T,C241T,C3037T,G28881A,G28882A,G28883C | N-G204R,N-R203K,ORF14-G50N,ORF1b-P314L,S-D614G |
| EPI_ISL_420599 | A23403G,C14408T,C18877T,C241T,C3037T,C5392T,C6449T,G25563T | ORF1a-L2062F,ORF1b-P314L,ORF3a-Q57H,S-D614G |
| EPI_ISL_420598 | A23403G,C14408T,C15324T,C241T,C28844T,C3037T,T10561C | N-R191C,ORF1b-P314L,S-D614G |
| EPI_ISL_430793 | A23403G,C14408T,C17165T,C241T,C3037T,G28881A,G28882A,G28883C,T27299C,T29148C | N-G204R,N-I292T,N-R203K,ORF14-G50N,ORF1b-P314L,ORF1b-S1233F,ORF6-I33T,S-D614G |
| EPI_ISL_430794 | A23403G,C12823T,C14408T,C241T,C24904T,C3037T,G25088T,G26727T,G28881A,G28882A,G28883C | M-A69S,N-G204R,N-R203K,ORF14-G50N,ORF1b-P314L,S-D614G,S-V1176F |
| EPI_ISL_430795 | A23403G,C10138T,C1059T,C14408T,C241T,C3037T,G15760A,G25563T | ORF1a-T265I,ORF1b-G765S,ORF1b-P314L,ORF3a-Q57H,S-D614G |
| EPI_ISL_430796 | A20268G,A23403G,C14408T,C241T,C25721G,C25904T,C3037T,G11083T | ORF1a-L3606F,ORF1b-P314L,ORF3a-A110G,ORF3a-S171L,S-D614G |
| EPI_ISL_430797 | A23403G,C14408T,C1473T,C241T,C3037T,G28881A,G28882A,G28883C,G29473A | N-G204R,N-R203K,ORF14-G50N,ORF1a-T403I,ORF1b-P314L,S-D614G |
| EPI_ISL_430798 | A23403G,C1059T,C11916T,C14408T,C18998T,C241T,C3037T,G25563T,G29540A | ORF1a-S3884L,ORF1a-T265I,ORF1b-A1844V,ORF1b-P314L,ORF3a-Q57H,S-D614G |
| EPI_ISL_430799 | A20268G,A23403G,C14408T,C241T,C3037T | ORF1b-P314L,S-D614G |
| EPI_ISL_430800 | A20268G,A23403G,C14408T,C241T,C3037T | ORF1b-P314L,S-D614G |
| EPI_ISL_430801 | A20268G,A23403G,C14408T,C241T,C3037T | ORF1b-P314L,S-D614G |
| EPI_ISL_430802 | A23403G,C14408T,C241T,C3037T,G28881A,G28882A,G28883C,T20989A | N-G204R,N-R203K,ORF14-G50N,ORF1b-L2508M,ORF1b-P314L,S-D614G |
| EPI_ISL_430803 | A20268G,A23403G,C12213T,C14408T,C22088T,C241T,C3037T | ORF1a-S3983F,ORF1b-P314L,S-D614G,S-L176F |
| EPI_ISL_430804 | A20268G,A23403G,C14408T,C241T,C3037T | ORF1b-P314L,S-D614G |
| EPI_ISL_430805 | A23403G,C14408T,C1473T,C16869T,C241T,C3037T,G28881A,G28882A,G28883C,G29473A | N-G204R,N-R203K,ORF14-G50N,ORF1a-T403I,ORF1b-P314L,S-D614G |
| EPI_ISL_430806 | A20268G,A23403G,C14408T,C241T,C3037T | ORF1b-P314L,S-D614G |
| EPI_ISL_430807 | A23403G,C14408T,C241T,C3037T,G28881A,G28882A,G28883C | N-G204R,N-R203K,ORF14-G50N,ORF1b-P314L,S-D614G |
| EPI_ISL_430808 | A23403G,C13536T,C14408T,C23731T,C241T,C3037T,C4002T,G10097A,G11083T,G25660T,G28881A,G28882A,G28883C | N-G204R,N-R203K,ORF14-G50N,ORF1a-G3278S,ORF1a-L3606F,ORF1a-T1246I,ORF1b-P314L,ORF3a-V90F,S-D614G |
| EPI_ISL_430809 | A23403G,C13536T,C14408T,C18246T,C23731T,C241T,C3037T,C4002T,G10097A,G11083T,G28881A,G28882A,G28883C | N-G204R,N-R203K,ORF14-G50N,ORF1a-G3278S,ORF1a-L3606F,ORF1a-T1246I,ORF1b-P314L,S-D614G |
| EPI_ISL_430810 | A23403G,C1059T,C11916T,C14362T,C14408T,C18998T,C241T,C3037T,G16852T,G25563T,G29540A | ORF1a-S3884L,ORF1a-T265I,ORF1b-A1844V,ORF1b-G1129C,ORF1b-P314L,ORF3a-Q57H,S-D614G |
| EPI_ISL_430811 | A23403G,C13536T,C14408T,C23731T,C241T,C3037T,C4002T,G10097A,G10157T,G28881A,G28882A,G28883C | N-G204R,N-R203K,ORF14-G50N,ORF1a-G3278S,ORF1a-T1246I,ORF1a-V3298L,ORF1b-P314L,S-D614G |
| EPI_ISL_430812 | A23403G,C13536T,C14408T,C23731T,C241T,C26801T,C3037T,C4002T,G10097A,G10157T,G25775T,G28881A,G28882A,G28883 | N-G204R,N-R203K,ORF14-G50N,ORF1a-G3278S,ORF1a-T1246I,ORF1a-V3298L,ORF1b-P314L,ORF3a-W128L,S-D614G |
| EPI_ISL_430813 | A23403G,C13536T,C14408T,C23731T,C241T,C3037T,C4002T,G10097A,G10157T,G28881A,G28882A,G28883C | N-G204R,N-R203K,ORF14-G50N,ORF1a-G3278S,ORF1a-T1246I,ORF1a-V3298L,ORF1b-P314L,S-D614G |
| EPI_ISL_430814 | A23403G,C14408T,C241T,C3037T,G26416T,G28881A,G28882A,G28883C,T27299C,T29148C | E-V58F,N-G204R,N-I292T,N-R203K,ORF14-G50N,ORF1b-P314L,ORF6-I33T,S-D614G |
| EPI_ISL_430815 | A23403G,C14408T,C241T,C26533T,C3037T,G26416T,G28881A,G28882A,G28883C,T27299C,T29148C | E-V58F,M-S4F,N-G204R,N-I292T,N-R203K,ORF14-G50N,ORF1b-P314L,ORF6-I33T,S-D614G |
| EPI_ISL_430816 | A23403G,C1059T,C11916T,C14408T,C18998T,C241T,C28863T,C3037T,G25563T,G29540A | N-S197L,ORF14-Q44*,ORF1a-S3884L,ORF1a-T265I,ORF1b-A1844V,ORF1b-P314L,ORF3a-Q57H,S-D614G |
| EPI_ISL_430817 | A23403G,C14408T,C241T,C3037T,G2173A,G26416T,G28881A,G28882A,G28883C,T27299C,T29148C | E-V58F,N-G204R,N-I292T,N-R203K,ORF14-G50N,ORF1b-P314L,ORF6-I33T,S-D614G |
| EPI_ISL_430818 | A23403G,A5608G,C14408T,C241T,C24789T,C26882T,C3037T,C313T,G20995A,G25522A,G28881A,G28882A,G28883C | N-G204R,N-R203K,ORF14-G50N,ORF1b-G2510S,ORF1b-P314L,ORF3a-G44R,S-D614G,S-T1076I |
| EPI_ISL_407893 | C8782T,T28144C | ORF8-L84S |
| EPI_ISL_408976 | Not_Found | Not_Found |
| EPI_ISL_408977 | Not_Found | Not_Found |
| EPI_ISL_417030 | C2445T,C29095T,C8782T,T20367C,T28144C | ORF1a-T727I,ORF8-L84S |
| EPI_ISL_412975 | A20047G,G11083T,G1397A,G29742T,G4255A,T28688C | ORF1a-L3606F,ORF1a-V378I,ORF1b-I2194V |
| EPI_ISL_413213 | C24904T,C884T,G11083T,G1397A,G29742T,G8653T,T28688C | ORF1a-L3606F,ORF1a-M2796I,ORF1a-R207C,ORF1a-V378I |
| EPI_ISL_413214 | G11083T,G1397A,G29374A,G29742T,T28688C | ORF1a-L3606F,ORF1a-V378I |
| EPI_ISL_413594 | G25323T | S-C1254F |
| EPI_ISL_413595 | C24381T,C29627T,C884T,G11083T,G1397A,G28457A,G29742T,G8653T,T28688C | N-E62K,ORF10-R24C,ORF1a-L3606F,ORF1a-M2796I,ORF1a-R207C,ORF1a-V378I,S-S940F |
| EPI_ISL_427722 | A17858G,C17747T,C18060T,C8782T,G29477T,T28144C | N-D402Y,ORF1b-P1427L,ORF1b-Y1464C,ORF8-L84S |
| EPI_ISL_427717 | C21005T,C2106T,C683T,T4057C | ORF1a-T614I,ORF1b-A2513V |
| EPI_ISL_413596 | G25323T | S-C1254F |
| EPI_ISL_427656 | A23403G,C14408T,C241T,C3037T | ORF1b-P314L,S-D614G |
| EPI_ISL_427657 | A23403G,C14408T,C241T,C3037T | ORF1b-P314L,S-D614G |
| EPI_ISL_427677 | A23403G,C14408T,C241T,C3037T | ORF1b-P314L,S-D614G |
| EPI_ISL_427686 | A23403G,C14408T,C241T,C3037T | ORF1b-P314L,S-D614G |
| EPI_ISL_427748 | A23403G,C14408T,C241T,C3037T | ORF1b-P314L,S-D614G |
| EPI_ISL_427772 | A23403G,C14408T,C241T,C3037T | ORF1b-P314L,S-D614G |
| EPI_ISL_427804 | A23403G,C14408T,C241T,C3037T | ORF1b-P314L,S-D614G |
| EPI_ISL_427754 | A20268G,A23403G,C14408T,C241T,C3037T | ORF1b-P314L,S-D614G |
| EPI_ISL_427741 | A20268G,A23403G,C14408T,C241T,C3037T | ORF1b-P314L,S-D614G |
| EPI_ISL_427661 | A20268G,A23403G,C14408T,C241T,C3037T,G25483A | ORF1b-P314L,ORF3a-A31T,S-D614G |
| EPI_ISL_413597 | C884T,G11083T,G1397A,G29742T,G8653T,T28688C | ORF1a-L3606F,ORF1a-M2796I,ORF1a-R207C,ORF1a-V378I |
| EPI_ISL_427666 | A20268G,A23403G,C14408T,C241T,C3037T,G25483A | ORF1b-P314L,ORF3a-A31T,S-D614G |
| EPI_ISL_427728 | A20268G,A23403G,C14408T,C241T,C3037T,G25483A | ORF1b-P314L,ORF3a-A31T,S-D614G |
| EPI_ISL_427724 | A20268G,A23403G,C14408T,C241T,C3037T,G17259A | ORF1b-P314L,S-D614G |
| EPI_ISL_427696 | A23403G,C14408T,C241T,C3037T,C6723T,T26094C | ORF1a-T2153I,ORF1b-P314L,S-D614G |
| EPI_ISL_427698 | A23403G,C14408T,C241T,C3037T,C6723T,G29227T,T26094C | ORF1a-T2153I,ORF1b-P314L,S-D614G |
| EPI_ISL_427756 | A20268G,A23403G,C14408T,C241T,C3037T,G15732T,T19143C | ORF1b-P314L,S-D614G |
| EPI_ISL_427693 | A15175G,A23403G,C14408T,C241T,C3037T,G29315T | N-D348Y,ORF1b-I570V,ORF1b-P314L,S-D614G |
| EPI_ISL_427669 | A23403G,C1059T,C10851T,C14408T,C241T,C3037T,G25563T | ORF1a-A3529V,ORF1a-T265I,ORF1b-P314L,ORF3a-Q57H,S-D614G |
| EPI_ISL_427679 | A23403G,C1059T,C10851T,C14408T,C241T,C3037T,G25563T | ORF1a-A3529V,ORF1a-T265I,ORF1b-P314L,ORF3a-Q57H,S-D614G |
| EPI_ISL_427688 | A23403G,C1059T,C10851T,C14408T,C241T,C3037T,G25563T | ORF1a-A3529V,ORF1a-T265I,ORF1b-P314L,ORF3a-Q57H,S-D614G |
| EPI_ISL_413598 | C24381T,C29627T,C884T,G11083T,G1397A,G28457A,G29742T,G8653T,T28688C | N-E62K,ORF10-R24C,ORF1a-L3606F,ORF1a-M2796I,ORF1a-R207C,ORF1a-V378I,S-S940F |
| EPI_ISL_427778 | A23403G,C1059T,C10851T,C14408T,C241T,C3037T,G25563T | ORF1a-A3529V,ORF1a-T265I,ORF1b-P314L,ORF3a-Q57H,S-D614G |
| EPI_ISL_427780 | A23403G,C1059T,C10851T,C14408T,C241T,C3037T,G25563T | ORF1a-A3529V,ORF1a-T265I,ORF1b-P314L,ORF3a-Q57H,S-D614G |
| EPI_ISL_427782 | A23403G,C1059T,C10851T,C14408T,C241T,C3037T,G25563T | ORF1a-A3529V,ORF1a-T265I,ORF1b-P314L,ORF3a-Q57H,S-D614G |
| EPI_ISL_427785 | A23403G,C1059T,C10851T,C14408T,C241T,C3037T,G25563T | ORF1a-A3529V,ORF1a-T265I,ORF1b-P314L,ORF3a-Q57H,S-D614G |
| EPI_ISL_427794 | A23403G,C1059T,C10851T,C14408T,C241T,C3037T,G25563T | ORF1a-A3529V,ORF1a-T265I,ORF1b-P314L,ORF3a-Q57H,S-D614G |
| EPI_ISL_427808 | A23403G,C1059T,C10851T,C14408T,C241T,C3037T,G25563T | ORF1a-A3529V,ORF1a-T265I,ORF1b-P314L,ORF3a-Q57H,S-D614G |
| EPI_ISL_427689 | A23403G,C1059T,C10851T,C14408T,C241T,C29370T,C3037T,G25563T | N-T366I,ORF1a-A3529V,ORF1a-T265I,ORF1b-P314L,ORF3a-Q57H,S-D614G |
| EPI_ISL_427760 | A23403G,C1059T,C10851T,C14408T,C241T,C3037T,C4890T,G25563T | ORF1a-A3529V,ORF1a-T1542I,ORF1a-T265I,ORF1b-P314L,ORF3a-Q57H,S-D614G |
| EPI_ISL_427768 | A23403G,C1059T,C10851T,C14408T,C1812T,C241T,C3037T,G25563T | ORF1a-A3529V,ORF1a-A516V,ORF1a-T265I,ORF1b-P314L,ORF3a-Q57H,S-D614G |
| EPI_ISL_427783 | A23403G,C1059T,C10851T,C14408T,C241T,C3037T,C5365T,G25563T | ORF1a-A3529V,ORF1a-T265I,ORF1b-P314L,ORF3a-Q57H,S-D614G |
| EPI_ISL_413599 | C18928T,C2113T,G11083T,G1397A,G29374A,G29742T,T28688C | ORF1a-L3606F,ORF1a-V378I,ORF1b-P1821S |
| EPI_ISL_427787 | A23403G,C1059T,C10851T,C14408T,C241T,C3037T,C6730T,G25563T | ORF1a-A3529V,ORF1a-T265I,ORF1b-P314L,ORF3a-Q57H,S-D614G |
| EPI_ISL_427791 | A23403G,C1059T,C10851T,C14408T,C18788T,C241T,C3037T,G25563T | ORF1a-A3529V,ORF1a-T265I,ORF1b-P314L,ORF1b-T1774I,ORF3a-Q57H,S-D614G |
| EPI_ISL_427742 | A23403G,C1059T,C10851T,C14408T,C15763T,C241T,C3037T,G25563T | ORF1a-A3529V,ORF1a-T265I,ORF1b-P314L,ORF3a-Q57H,S-D614G |
| EPI_ISL_427673 | A23403G,C1059T,C12885T,C14408T,C241T,C3037T,G25563T | ORF1a-T265I,ORF1a-T4207I,ORF1b-P314L,ORF3a-Q57H,S-D614G |
| EPI_ISL_427692 | A23403G,C1059T,C14408T,C241T,C3037T,G25563T | ORF1a-T265I,ORF1b-P314L,ORF3a-Q57H,S-D614G |
| EPI_ISL_427700 | A23403G,C1059T,C14408T,C241T,C3037T,G25563T | ORF1a-T265I,ORF1b-P314L,ORF3a-Q57H,S-D614G |
| EPI_ISL_427761 | A23403G,C1059T,C14408T,C241T,C3037T,G25563T | ORF1a-T265I,ORF1b-P314L,ORF3a-Q57H,S-D614G |
| EPI_ISL_427795 | A23403G,C10138T,C1059T,C14408T,C241T,C3037T,G25563T | ORF1a-T265I,ORF1b-P314L,ORF3a-Q57H,S-D614G |
| EPI_ISL_427799 | A23403G,C10138T,C1059T,C14408T,C241T,C3037T,G25563T | ORF1a-T265I,ORF1b-P314L,ORF3a-Q57H,S-D614G |
| EPI_ISL_427802 | A23403G,C1059T,C14408T,C241T,C3037T,G13822A,G25563T | ORF1a-T265I,ORF1b-P314L,ORF1b-V119I,ORF3a-Q57H,S-D614G |
| EPI_ISL_413600 | A20047G,G11083T,G1397A,G29742T,G4255A,T28688C | ORF1a-L3606F,ORF1a-V378I,ORF1b-I2194V |
| EPI_ISL_427749 | A23403G,C1059T,C13501T,C14408T,C241T,C27128T,C3037T,G25563T | ORF1a-T265I,ORF1b-P12S,ORF1b-P314L,ORF3a-Q57H,S-D614G |
| EPI_ISL_427737 | A23403G,A6749G,C1059T,C14408T,C241T,C3037T,G15380T,G25563T | ORF1a-N2162D,ORF1a-T265I,ORF1b-P314L,ORF1b-S638I,ORF3a-Q57H,S-D614G |
| EPI_ISL_427738 | A23403G,C1059T,C14408T,C241T,C3037T,G15380T,G25563T | ORF1a-T265I,ORF1b-P314L,ORF1b-S638I,ORF3a-Q57H,S-D614G |
| EPI_ISL_427708 | A23403G,C1059T,C10851T,C14408T,C241T,C3037T,G14122T,G25563T | ORF1a-A3529V,ORF1a-T265I,ORF1b-G219C,ORF1b-P314L,ORF3a-Q57H,S-D614G |
| EPI_ISL_427682 | A23403G,C1059T,C10851T,C14408T,C23439T,C241T,C3037T,G20931T,G25563T | ORF1a-A3529V,ORF1a-T265I,ORF1b-P314L,ORF3a-Q57H,S-A626V,S-D614G |
| EPI_ISL_427764 | A23403G,C1059T,C10851T,C14408T,C23439T,C241T,C3037T,G20931T,G25563T | ORF1a-A3529V,ORF1a-T265I,ORF1b-P314L,ORF3a-Q57H,S-A626V,S-D614G |
| EPI_ISL_427683 | A23403G,C1059T,C10851T,C14408T,C17825T,C23439T,C241T,C3037T,G20931T,G25563T | ORF1a-A3529V,ORF1a-T265I,ORF1b-P314L,ORF1b-T1453I,ORF3a-Q57H,S-A626V,S-D614G |
| EPI_ISL_427775 | A23403G,C1059T,C10851T,C14408T,C23439T,C241T,C3037T,G20931T,G25563T,G7787T | ORF1a-A2508S,ORF1a-A3529V,ORF1a-T265I,ORF1b-P314L,ORF3a-Q57H,S-A626V,S-D614G |
| EPI_ISL_427796 | A23403G,C14408T,C241T,C26461T,C3037T,C7083T,G25429T | E-L73F,ORF1a-S2273F,ORF1b-P314L,ORF3a-V13L,S-D614G |
| EPI_ISL_427721 | A23403G,C1059T,C14408T,C21642T,C241T,C25688T,C3037T,G25563T,T13006C,T18768C | ORF1a-T265I,ORF1b-P314L,ORF3a-A99V,ORF3a-Q57H,S-A27V,S-D614G |
