## Supplementary Table 2 for "What if we perceive SARS-CoV-2 genomes as documents? Topic modelling using Latent Dirichlet Allocation to identify mutation signatures and classify SARS-CoV-2 genomes"

**Supplementary Table 2**: Bi-grams detected in Nucleotide mutations with a minimum co-occurence in 500 genomes

| Bi-gram | Score | Count |
|---|---|---|
| G28882A,G28883C | 4.1696214807 | 13413 |
| G28881A,G28882A | 4.1572109611 | 13412 |
| A23403G,G25563T | 1.362210351 | 9282 |
| C241T,C1059T | 1.4286804332 | 7643 |
| C1059T,C3037T | 1.3228613655 | 7098 |
| G11083T,C14805T | 5.5556961849 | 3050 |
| A20268G,A23403G | 1.0517649495 | 1980 |
| A2480G,C2558T | 22.4309194799 | 1721 |
| A17858G,C18060T | 23.0890879075 | 1708 |
| C17747T,A17858G | 23.3715377437 | 1688 |
| A23403G,C23731T | 1.1168434932 | 1676 |
| C14805T,G26144T | 2.2677768664 | 1235 |
| C14805T,T17247C | 6.5630809005 | 1191 |
| C8782T,C17747T | 5.4912759033 | 1097 |
| C2558T,G11083T | 3.046028963 | 1044 |
| G1440A,G2891A | 24.7702260936 | 951 |
| C18060T,T28144C | 3.8552140865 | 937 |
| T17247C,G26144T | 4.2750995704 | 884 |
| C8782T,T9477A | 6.1218867196 | 832 |
| C28657T,C28863T | 25.3936598004 | 830 |
| G25563T,C27964T | 1.3851741982 | 733 |
| T9477A,C14805T | 3.3383274673 | 730 |
| G28580T,G28881A | 1.2757541217 | 718 |
| T28144C,C28657T | 3.1475336353 | 679 |
| G25979T,T28144C | 2.8815680315 | 657 |
| C23731T,G28580T | 1.9747877866 | 544 |
| C4002T,G10097A | 1.08036905 | 521 |
