## Supplementary Table 3 for "What if we perceive SARS-CoV-2 genomes as documents? Topic modelling using Latent Dirichlet Allocation to identify mutation signatures and classify SARS-CoV-2 genomes"

**Supplementary Table 3:** Tri-grams detected in Nucleotide mutations with a minimum co-occurence in 500 genomes

| Tri-grams | Score | Count |
|---|---|---|
| G28881A_G28882A_G28883C | 26721.5651537335 | 12730 |
| A23403G_G28881A_G28882A | 1.3737656473 | 6928 |
| C241T_C1059T_C3037T | 24.5000362592 | 6556 |
| C14408T_A23403G_G25563T | 17.5480691782 | 5183 |
| C14408T_A20268G_A23403G | 10.0180266462 | 1461 |
| C14408T_A23403G_C23731T | 91.9008184102 | 1241 |
| G11083T_C14805T_G26144T | 58.091512809 | 1028 |
| C18877T_A23403G_G25563T | 1.2983430221 | 735 |
| C8782T_T9477A_C14805T | 422.7844245669 | 713 |
| A23403G_G25563T_C27964T | 3.6186863576 | 633 |
| A17858G_C18060T_T28144C | 4665.1939736347 | 583 |
| C3037T_C4002T_G10097A | 2.9868614808 | 507 |
