## Supplementary Table 4 for "What if we perceive SARS-CoV-2 genomes as documents? Topic modelling using Latent Dirichlet Allocation to identify mutation signatures and classify SARS-CoV-2 genomes"

| Gisaid_EPI_I | Originating_Lab | Submitting_Lab | Authors | Title | Paper_URL |
|---|---|---|---|---|---|
| EPI_ISL_418241 | NIC Viral Respiratory Unit - Institut Pasteur of Algeria | National Reference Center for Viruses of Respiratory Infections, Institut Pasteur, Paris | Mélanie Albert et al | ? | ? |
| EPI_ISL_418242 | NIC Viral Respiratory Unit - Institut Pasteur of Algeria | National Reference Center for Viruses of Respiratory Infections, Institut Pasteur, Paris | Mélanie Albert et al | ? | ? |
| EPI_ISL_420037 | NIC Viral Respiratory Unit - Institut Pasteur of Algeria | National Reference Center for Viruses of Respiratory Infections, Institut Pasteur, Paris | Mélanie Albert et al | ? | ? |
| EPI_ISL_452463 | Hospital Universitario Puerta del Mar de Cádiz - INIBICA | SeqCOVID-SPAIN consortium_IBV(CSIC) | Salud Rodríguez-Pallares et al | ? | ? |
| EPI_ISL_413485 | Department of microbiology laboratory,Anhui Provincial Center for Disease Control and Prevention | Department of microbiology laboratory,Anhui Provincial Center for Disease Control and Prevention | Weiwei Li et al | ? | ? |
| EPI_ISL_420600 | Servicio Virosis Respiratorias-Departamento Virología-INEI | Instituto Nacional Enfermedades Infecciosas C.G.Malbran | Baumeister E. et al | COVID-19 in Latin America: Contrasting phylodynamic inference with epidemiological surveillance. | https://dx.doi.org_10.1101_2020.05.23.20111443 |
| EPI_ISL_420599 | Servicio Virosis Respiratorias-Departamento Virología-INEI | Instituto Nacional Enfermedades Infecciosas C.G.Malbran | Baumeister E. et al | COVID-19 in Latin America: Contrasting phylodynamic inference with epidemiological surveillance. | https://dx.doi.org_10.1101_2020.05.23.20111443 |
| EPI_ISL_420598 | Servicio Virosis Respiratorias-Departamento Virología-INEI | Instituto Nacional Enfermedades Infecciosas C.G.Malbran | Baumeister E. et al | COVID-19 in Latin America: Contrasting phylodynamic inference with epidemiological surveillance. | https://dx.doi.org_10.1101_2020.05.23.20111443 |
| EPI_ISL_430793 | Laboratorio Análisis Clínicos, Unidad de Servicios Diagnósticos, Swiss Medical Group | Área de Secuenciación del Laboratorio de Virología del Hospital de Niños Dr. Ricardo Gutierrez | Nabaes Jodar et al | COVID-19 in Latin America: Contrasting phylodynamic inference with epidemiological surveillance. | https://dx.doi.org_10.1101_2020.05.23.20111443 |
| EPI_ISL_430794 | Laboratorio Análisis Clínicos, Unidad de Servicios Diagnósticos, Swiss Medical Group | Área de Secuenciación del Laboratorio de Virología del Hospital de Niños Dr. Ricardo Gutierrez | Nabaes Jodar et al | ? | ? |
| EPI_ISL_430795 | Laboratorio de Virología del Hospital de Niños Dr. Ricardo Gutierrez | Área de Secuenciación del Laboratorio de Virología del Hospital de Niños Dr. Ricardo Gutierrez | Nabaes Jodar et al | ? | ? |
| EPI_ISL_430796 | Departamento de Biología y genética molecular, IACA Laboratorios. | Área de Secuenciación del Laboratorio de Virología del Hospital de Niños Dr. Ricardo Gutierrez | Nabaes Jodar et al | COVID-19 in Latin America: Contrasting phylodynamic inference with epidemiological surveillance. | https://dx.doi.org_10.1101_2020.05.23.20111443 |
| EPI_ISL_430797 | Departamento de Biología y genética molecular, IACA Laboratorios. | Área de Secuenciación del Laboratorio de Virología del Hospital de Niños Dr. Ricardo Gutierrez | Nabaes Jodar et al | COVID-19 in Latin America: Contrasting phylodynamic inference with epidemiological surveillance. | https://dx.doi.org_10.1101_2020.05.23.20111443 |
| EPI_ISL_430798 | Departamento de Biología y genética molecular, IACA Laboratorios. | Área de Secuenciación del Laboratorio de Virología del Hospital de Niños Dr. Ricardo Gutierrez | Nabaes Jodar et al | COVID-19 in Latin America: Contrasting phylodynamic inference with epidemiological surveillance. | https://dx.doi.org_10.1101_2020.05.23.20111443 |
| EPI_ISL_430799 | Laboratorio de Virología del Hospital de Niños Dr. Ricardo Gutierrez | Área de Secuenciación del Laboratorio de Virología del Hospital de Niños Dr. Ricardo Gutierrez | Nabaes Jodar et al | COVID-19 in Latin America: Contrasting phylodynamic inference with epidemiological surveillance. | https://dx.doi.org_10.1101_2020.05.23.20111443 |
| EPI_ISL_430800 | Laboratorio de Virología del Hospital de Niños Dr. Ricardo Gutierrez | Área de Secuenciación del Laboratorio de Virología del Hospital de Niños Dr. Ricardo Gutierrez | Nabaes Jodar et al | ? | ? |
| EPI_ISL_430801 | Laboratorio de Virología del Hospital de Niños Dr. Ricardo Gutierrez | Área de Secuenciación del Laboratorio de Virología del Hospital de Niños Dr. Ricardo Gutierrez | Nabaes Jodar et al | COVID-19 in Latin America: Contrasting phylodynamic inference with epidemiological surveillance. | https://dx.doi.org_10.1101_2020.05.23.20111443 |
| EPI_ISL_430802 | Departamento de Biología y genética molecular, IACA Laboratorios. | Área de Secuenciación del Laboratorio de Virología del Hospital de Niños Dr. Ricardo Gutierrez | Nabaes Jodar et al | COVID-19 in Latin America: Contrasting phylodynamic inference with epidemiological surveillance. | https://dx.doi.org_10.1101_2020.05.23.20111443 |
| EPI_ISL_430803 | Laboratorio de Virología del Hospital de Niños Dr. Ricardo Gutierrez | Área de Secuenciación del Laboratorio de Virología del Hospital de Niños Dr. Ricardo Gutierrez | Nabaes Jodar et al | COVID-19 in Latin America: Contrasting phylodynamic inference with epidemiological surveillance. | https://dx.doi.org_10.1101_2020.05.23.20111443 |
| EPI_ISL_430804 | Laboratorio de Virología del Hospital de Niños Dr. Ricardo Gutierrez | Área de Secuenciación del Laboratorio de Virología del Hospital de Niños Dr. Ricardo Gutierrez | Nabaes Jodar et al | COVID-19 in Latin America: Contrasting phylodynamic inference with epidemiological surveillance. | https://dx.doi.org_10.1101_2020.05.23.20111443 |
| EPI_ISL_430805 | Departamento de Biología y genética molecular, IACA Laboratorios. | Área de Secuenciación del Laboratorio de Virología del Hospital de Niños Dr. Ricardo Gutierrez | Nabaes Jodar et al | COVID-19 in Latin America: Contrasting phylodynamic inference with epidemiological surveillance. | https://dx.doi.org_10.1101_2020.05.23.20111443 |
| EPI_ISL_430806 | Departamento de Biología y genética molecular, IACA Laboratorios. | Área de Secuenciación del Laboratorio de Virología del Hospital de Niños Dr. Ricardo Gutierrez | Nabaes Jodar et al | ? | ? |
| EPI_ISL_430807 | Laboratorio de Virología del Hospital de Niños Dr. Ricardo Gutierrez | Área de Secuenciación del Laboratorio de Virología del Hospital de Niños Dr. Ricardo Gutierrez | Nabaes Jodar et al | COVID-19 in Latin America: Contrasting phylodynamic inference with epidemiological surveillance. | https://dx.doi.org_10.1101_2020.05.23.20111443 |
| EPI_ISL_430808 | Departamento de Biología y genética molecular, IACA Laboratorios. | Área de Secuenciación del Laboratorio de Virología del Hospital de Niños Dr. Ricardo Gutierrez | Nabaes Jodar et al | COVID-19 in Latin America: Contrasting phylodynamic inference with epidemiological surveillance. | https://dx.doi.org_10.1101_2020.05.23.20111443 |
| EPI_ISL_430809 | Laboratorio de Virología del Hospital de Niños Dr. Ricardo Gutierrez | Área de Secuenciación del Laboratorio de Virología del Hospital de Niños Dr. Ricardo Gutierrez | Nabaes Jodar et al | COVID-19 in Latin America: Contrasting phylodynamic inference with epidemiological surveillance. | https://dx.doi.org_10.1101_2020.05.23.20111443 |
| EPI_ISL_430810 | Laboratorio de Virología del Hospital de Niños Dr. Ricardo Gutierrez | Área de Secuenciación del Laboratorio de Virología del Hospital de Niños Dr. Ricardo Gutierrez | Nabaes Jodar et al | COVID-19 in Latin America: Contrasting phylodynamic inference with epidemiological surveillance. | https://dx.doi.org_10.1101_2020.05.23.20111443 |
| EPI_ISL_430811 | Laboratorio de Virología del Hospital de Niños Dr. Ricardo Gutierrez | Área de Secuenciación del Laboratorio de Virología del Hospital de Niños Dr. Ricardo Gutierrez | Nabaes Jodar et al | COVID-19 in Latin America: Contrasting phylodynamic inference with epidemiological surveillance. | https://dx.doi.org_10.1101_2020.05.23.20111443 |
| EPI_ISL_430812 | Laboratorio de Virología del Hospital de Niños Dr. Ricardo Gutierrez | Área de Secuenciación del Laboratorio de Virología del Hospital de Niños Dr. Ricardo Gutierrez | Nabaes Jodar et al | COVID-19 in Latin America: Contrasting phylodynamic inference with epidemiological surveillance. | https://dx.doi.org_10.1101_2020.05.23.20111443 |
| EPI_ISL_430813 | Laboratorio de Virología del Hospital de Niños Dr. Ricardo Gutierrez | Área de Secuenciación del Laboratorio de Virología del Hospital de Niños Dr. Ricardo Gutierrez | Nabaes Jodar et al | ? | ? |
| EPI_ISL_430814 | Laboratorio de Virología del Hospital de Niños Dr. Ricardo Gutierrez | Área de Secuenciación del Laboratorio de Virología del Hospital de Niños Dr. Ricardo Gutierrez | Nabaes Jodar et al | COVID-19 in Latin America: Contrasting phylodynamic inference with epidemiological surveillance. | https://dx.doi.org_10.1101_2020.05.23.20111443 |
| EPI_ISL_430815 | Laboratorio de Virología del Hospital de Niños Dr. Ricardo Gutierrez | Área de Secuenciación del Laboratorio de Virología del Hospital de Niños Dr. Ricardo Gutierrez | Nabaes Jodar et al | COVID-19 in Latin America: Contrasting phylodynamic inference with epidemiological surveillance. | https://dx.doi.org_10.1101_2020.05.23.20111443 |
| EPI_ISL_430816 | Laboratorio de Virología del Hospital de Niños Dr. Ricardo Gutierrez | Área de Secuenciación del Laboratorio de Virología del Hospital de Niños Dr. Ricardo Gutierrez | Nabaes Jodar et al | COVID-19 in Latin America: Contrasting phylodynamic inference with epidemiological surveillance. | https://dx.doi.org_10.1101_2020.05.23.20111443 |
| EPI_ISL_430817 | Laboratorio de Virología del Hospital de Niños Dr. Ricardo Gutierrez | Área de Secuenciación del Laboratorio de Virología del Hospital de Niños Dr. Ricardo Gutierrez | Nabaes Jodar et al | COVID-19 in Latin America: Contrasting phylodynamic inference with epidemiological surveillance. | https://dx.doi.org_10.1101_2020.05.23.20111443 |
| EPI_ISL_430818 | Laboratorio de Virología del Hospital de Niños Dr. Ricardo Gutierrez | Área de Secuenciación del Laboratorio de Virología del Hospital de Niños Dr. Ricardo Gutierrez | Nabaes Jodar et al | COVID-19 in Latin America: Contrasting phylodynamic inference with epidemiological surveillance. | https://dx.doi.org_10.1101_2020.05.23.20111443 |
| EPI_ISL_407893 | Centre for Infectious Diseases and Microbiology Laboratory Services | NSW Health Pathology - Institute of Clinical Pathology and Medical Research; Westmead Hospital; University of Sydney | Eden J-S et al | Revealing COVID-19 Transmission by SARS-CoV-2 Genome Sequencing and Agent Based Modellin | https://dx.doi.org_10.1101_2020.04.19.048751 |
| EPI_ISL_408976 | Centre for Infectious Diseases and Microbiology Laboratory Services | NSW Health Pathology - Institute of Clinical Pathology and Medical Research; Westmead Hospital; University of Sydney | Rockett R et al | Revealing COVID-19 Transmission by SARS-CoV-2 Genome Sequencing and Agent Based Modellin | https://dx.doi.org_10.1101_2020.04.19.048751 |
| EPI_ISL_408977 | Serology, Virology and OTDS Laboratories (SAViD), NSW Health Pathology Randwick | NSW Health Pathology - Institute of Clinical Pathology and Medical Research; Centre for Infectious Diseases and Microbiology Laboratory Services; Westmead Hospital; University of Sydney | Eden J-S et al | Revealing COVID-19 Transmission by SARS-CoV-2 Genome Sequencing and Agent Based Modellin | https://dx.doi.org_10.1101_2020.04.19.048751 |
| EPI_ISL_417030 | Centre for Infectious Diseases and Microbiology Laboratory Services | NSW Health Pathology - Institute of Clinical Pathology and Medical Research; Westmead Hospital; University of Sydney | Eden J-S et al | Revealing COVID-19 Transmission by SARS-CoV-2 Genome Sequencing and Agent Based Modellin | https://dx.doi.org_10.1101_2020.04.19.048751 |
| EPI_ISL_412975 | Centre for Infectious Diseases and Microbiology Laboratory Services | NSW Health Pathology - Institute of Clinical Pathology and Medical Research; Westmead Hospital; University of Sydney | Eden J-S et al | Revealing COVID-19 Transmission by SARS-CoV-2 Genome Sequencing and Agent Based Modellin | https://dx.doi.org_10.1101_2020.04.19.048751 |
| EPI_ISL_413213 | Centre for Infectious Diseases and Microbiology Laboratory Services | NSW Health Pathology - Institute of Clinical Pathology and Medical Research; Westmead Hospital; University of Sydney | Eden J-S et al | Revealing COVID-19 Transmission by SARS-CoV-2 Genome Sequencing and Agent Based Modellin | https://dx.doi.org_10.1101_2020.04.19.048751 |
| EPI_ISL_413214 | Centre for Infectious Diseases and Microbiology Laboratory Services | NSW Health Pathology - Institute of Clinical Pathology and Medical Research; Westmead Hospital; University of Sydney | Eden J-S et al | Revealing COVID-19 Transmission by SARS-CoV-2 Genome Sequencing and Agent Based Modellin | https://dx.doi.org_10.1101_2020.04.19.048751 |
| EPI_ISL_413594 | Centre for Infectious Diseases and Microbiology Laboratory Services | NSW Health Pathology - Institute of Clinical Pathology and Medical Research; Westmead Hospital; University of Sydney | Rockett R et al | Revealing COVID-19 Transmission by SARS-CoV-2 Genome Sequencing and Agent Based Modellin | https://dx.doi.org_10.1101_2020.04.19.048751 |
| EPI_ISL_413595 | Centre for Infectious Diseases and Microbiology Laboratory Services | NSW Health Pathology - Institute of Clinical Pathology and Medical Research; Westmead Hospital; University of Sydney | Rockett R et al | Revealing COVID-19 Transmission by SARS-CoV-2 Genome Sequencing and Agent Based Modellin | https://dx.doi.org_10.1101_2020.04.19.048751 |
| EPI_ISL_427722 | Centre for Infectious Diseases and Microbiology Public Health | NSW Health Pathology - Institute of Clinical Pathology and Medical Research; Westmead Hospital; University of Sydney | Draper J et al | Revealing COVID-19 Transmission by SARS-CoV-2 Genome Sequencing and Agent Based Modellin | https://dx.doi.org_10.1101_2020.04.19.048751 |
| EPI_ISL_427717 | ACT Pathology, The Canberra Hospital | NSW Health Pathology - Institute of Clinical Pathology and Medical Research; Westmead Hospital; University of Sydney | Sim E et al | Revealing COVID-19 Transmission by SARS-CoV-2 Genome Sequencing and Agent Based Modellin | https://dx.doi.org_10.1101_2020.04.19.048751 |
| EPI_ISL_413596 | Centre for Infectious Diseases and Microbiology - Public Health | NSW Health Pathology - Institute of Clinical Pathology and Medical Research; Westmead Hospital; University of Sydney | Rockett R et al | Revealing COVID-19 Transmission by SARS-CoV-2 Genome Sequencing and Agent Based Modellin | https://dx.doi.org_10.1101_2020.04.19.048751 |
| EPI_ISL_427656 | Centre for Infectious Diseases and Microbiology Public Health | NSW Health Pathology - Institute of Clinical Pathology and Medical Research; Westmead Hospital; University of Sydney | Bachmann N et al | Revealing COVID-19 Transmission by SARS-CoV-2 Genome Sequencing and Agent Based Modellin | https://dx.doi.org_10.1101_2020.04.19.048751 |
| EPI_ISL_427657 | Centre for Infectious Diseases and Microbiology Public Health | NSW Health Pathology - Institute of Clinical Pathology and Medical Research; Westmead Hospital; University of Sydney | Rockett R et al | Revealing COVID-19 Transmission by SARS-CoV-2 Genome Sequencing and Agent Based Modellin | https://dx.doi.org_10.1101_2020.04.19.048751 |
| EPI_ISL_427677 | Centre for Infectious Diseases and Microbiology Public Health | NSW Health Pathology - Institute of Clinical Pathology and Medical Research; Westmead Hospital; University of Sydney | Lam C et al | Revealing COVID-19 Transmission by SARS-CoV-2 Genome Sequencing and Agent Based Modellin | https://dx.doi.org_10.1101_2020.04.19.048751 |
| EPI_ISL_427686 | Centre for Infectious Diseases and Microbiology Public Health | NSW Health Pathology - Institute of Clinical Pathology and Medical Research; Westmead Hospital; University of Sydney | Gray K et al | Revealing COVID-19 Transmission by SARS-CoV-2 Genome Sequencing and Agent Based Modellin | https://dx.doi.org_10.1101_2020.04.19.048751 |
| EPI_ISL_427748 | Centre for Infectious Diseases and Microbiology Public Health | NSW Health Pathology - Institute of Clinical Pathology and Medical Research; Westmead Hospital; University of Sydney | Timms V et al | Revealing COVID-19 Transmission by SARS-CoV-2 Genome Sequencing and Agent Based Modellin | https://dx.doi.org_10.1101_2020.04.19.048751 |
| EPI_ISL_427772 | Centre for Infectious Diseases and Microbiology Public Health | NSW Health Pathology - Institute of Clinical Pathology and Medical Research; Westmead Hospital; University of Sydney | Gall M et al | Revealing COVID-19 Transmission by SARS-CoV-2 Genome Sequencing and Agent Based Modellin | https://dx.doi.org_10.1101_2020.04.19.048751 |
| EPI_ISL_427804 | Centre for Infectious Diseases and Microbiology Public Health | NSW Health Pathology - Institute of Clinical Pathology and Medical Research; Westmead Hospital; University of Sydney | Arnott A et al | Revealing COVID-19 Transmission by SARS-CoV-2 Genome Sequencing and Agent Based Modellin | https://dx.doi.org_10.1101_2020.04.19.048751 |
| EPI_ISL_427754 | Centre for Infectious Diseases and Microbiology Public Health | NSW Health Pathology - Institute of Clinical Pathology and Medical Research; Westmead Hospital; University of Sydney | Sadsad R et al | Revealing COVID-19 Transmission by SARS-CoV-2 Genome Sequencing and Agent Based Modellin | https://dx.doi.org_10.1101_2020.04.19.048751 |
| EPI_ISL_427741 | Centre for Infectious Diseases and Microbiology Public Health | NSW Health Pathology - Institute of Clinical Pathology and Medical Research; Westmead Hospital; University of Sydney | Draper J et al | Revealing COVID-19 Transmission by SARS-CoV-2 Genome Sequencing and Agent Based Modellin | https://dx.doi.org_10.1101_2020.04.19.048751 |
| EPI_ISL_427661 | Centre for Infectious Diseases and Microbiology Public Health | NSW Health Pathology - Institute of Clinical Pathology and Medical Research; Westmead Hospital; University of Sydney | Sim E et al | Revealing COVID-19 Transmission by SARS-CoV-2 Genome Sequencing and Agent Based Modellin | https://dx.doi.org_10.1101_2020.04.19.048751 |
| EPI_ISL_413597 | Centre for Infectious Diseases and Microbiology- Public Health | NSW Health Pathology - Institute of Clinical Pathology and Medical Research; Westmead Hospital; University of Sydney | Lam C et al | Revealing COVID-19 Transmission by SARS-CoV-2 Genome Sequencing and Agent Based Modellin | https://dx.doi.org_10.1101_2020.04.19.048751 |
| EPI_ISL_427666 | Centre for Infectious Diseases and Microbiology Public Health | NSW Health Pathology - Institute of Clinical Pathology and Medical Research; Westmead Hospital; University of Sydney | Bachmann N et al | Revealing COVID-19 Transmission by SARS-CoV-2 Genome Sequencing and Agent Based Modellin | https://dx.doi.org_10.1101_2020.04.19.048751 |
| EPI_ISL_427728 | Centre for Infectious Diseases and Microbiology Public Health | NSW Health Pathology - Institute of Clinical Pathology and Medical Research; Westmead Hospital; University of Sydney | Rockett R et al | Revealing COVID-19 Transmission by SARS-CoV-2 Genome Sequencing and Agent Based Modellin | https://dx.doi.org_10.1101_2020.04.19.048751 |
| EPI_ISL_427724 | Centre for Infectious Diseases and Microbiology Public Health | NSW Health Pathology - Institute of Clinical Pathology and Medical Research; Westmead Hospital; University of Sydney | Lam C et al | Revealing COVID-19 Transmission by SARS-CoV-2 Genome Sequencing and Agent Based Modellin | https://dx.doi.org_10.1101_2020.04.19.048751 |
| EPI_ISL_427696 | Centre for Infectious Diseases and Microbiology Public Health | NSW Health Pathology - Institute of Clinical Pathology and Medical Research; Westmead Hospital; University of Sydney | Gray K et al | Revealing COVID-19 Transmission by SARS-CoV-2 Genome Sequencing and Agent Based Modellin | https://dx.doi.org_10.1101_2020.04.19.048751 |
| EPI_ISL_427698 | Centre for Infectious Diseases and Microbiology Public Health | NSW Health Pathology - Institute of Clinical Pathology and Medical Research; Westmead Hospital; University of Sydney | Timms V et al | Revealing COVID-19 Transmission by SARS-CoV-2 Genome Sequencing and Agent Based Modellin | https://dx.doi.org_10.1101_2020.04.19.048751 |
| EPI_ISL_427756 | Centre for Infectious Diseases and Microbiology Public Health | NSW Health Pathology - Institute of Clinical Pathology and Medical Research; Westmead Hospital; University of Sydney | Gall M et al | Revealing COVID-19 Transmission by SARS-CoV-2 Genome Sequencing and Agent Based Modellin | https://dx.doi.org_10.1101_2020.04.19.048751 |
| EPI_ISL_427693 | Centre for Infectious Diseases and Microbiology Public Health | NSW Health Pathology - Institute of Clinical Pathology and Medical Research; Westmead Hospital; University of Sydney | Arnott A et al | Revealing COVID-19 Transmission by SARS-CoV-2 Genome Sequencing and Agent Based Modellin | https://dx.doi.org_10.1101_2020.04.19.048751 |
| EPI_ISL_427669 | Centre for Infectious Diseases and Microbiology Public Health | NSW Health Pathology - Institute of Clinical Pathology and Medical Research; Westmead Hospital; University of Sydney | Sadsad R et al | Revealing COVID-19 Transmission by SARS-CoV-2 Genome Sequencing and Agent Based Modellin | https://dx.doi.org_10.1101_2020.04.19.048751 |
| EPI_ISL_427679 | Centre for Infectious Diseases and Microbiology Public Health | NSW Health Pathology - Institute of Clinical Pathology and Medical Research; Westmead Hospital; University of Sydney | Draper J et al | Revealing COVID-19 Transmission by SARS-CoV-2 Genome Sequencing and Agent Based Modellin | https://dx.doi.org_10.1101_2020.04.19.048751 |
| EPI_ISL_427688 | Centre for Infectious Diseases and Microbiology Public Health | NSW Health Pathology - Institute of Clinical Pathology and Medical Research; Westmead Hospital; University of Sydney | Sim E et al | Revealing COVID-19 Transmission by SARS-CoV-2 Genome Sequencing and Agent Based Modellin | https://dx.doi.org_10.1101_2020.04.19.048751 |
| EPI_ISL_413598 | Centre for Infectious Diseases and Microbiology - Public Health | NSW Health Pathology - Institute of Clinical Pathology and Medical Research; Westmead Hospital; University of Sydney | Gray K et al | Revealing COVID-19 Transmission by SARS-CoV-2 Genome Sequencing and Agent Based Modellin | https://dx.doi.org_10.1101_2020.04.19.048751 |
| EPI_ISL_427778 | Centre for Infectious Diseases and Microbiology Public Health | NSW Health Pathology - Institute of Clinical Pathology and Medical Research; Westmead Hospital; University of Sydney | Bachmann N et al | Revealing COVID-19 Transmission by SARS-CoV-2 Genome Sequencing and Agent Based Modellin | https://dx.doi.org_10.1101_2020.04.19.048751 |
| EPI_ISL_427780 | Centre for Infectious Diseases and Microbiology Public Health | NSW Health Pathology - Institute of Clinical Pathology and Medical Research; Westmead Hospital; University of Sydney | Rockett R et al | Revealing COVID-19 Transmission by SARS-CoV-2 Genome Sequencing and Agent Based Modellin | https://dx.doi.org_10.1101_2020.04.19.048751 |
| EPI_ISL_427782 | Centre for Infectious Diseases and Microbiology Public Health | NSW Health Pathology - Institute of Clinical Pathology and Medical Research; Westmead Hospital; University of Sydney | Lam C et al | Revealing COVID-19 Transmission by SARS-CoV-2 Genome Sequencing and Agent Based Modellin | https://dx.doi.org_10.1101_2020.04.19.048751 |
| EPI_ISL_427785 | Centre for Infectious Diseases and Microbiology Public Health | NSW Health Pathology - Institute of Clinical Pathology and Medical Research; Westmead Hospital; University of Sydney | Gray K et al | Revealing COVID-19 Transmission by SARS-CoV-2 Genome Sequencing and Agent Based Modellin | https://dx.doi.org_10.1101_2020.04.19.048751 |
| EPI_ISL_427794 | Centre for Infectious Diseases and Microbiology Public Health | NSW Health Pathology - Institute of Clinical Pathology and Medical Research; Westmead Hospital; University of Sydney | Timms V et al | Revealing COVID-19 Transmission by SARS-CoV-2 Genome Sequencing and Agent Based Modellin | https://dx.doi.org_10.1101_2020.04.19.048751 |
| EPI_ISL_427808 | Centre for Infectious Diseases and Microbiology Public Health | NSW Health Pathology - Institute of Clinical Pathology and Medical Research; Westmead Hospital; University of Sydney | Gall M et al | Revealing COVID-19 Transmission by SARS-CoV-2 Genome Sequencing and Agent Based Modellin | https://dx.doi.org_10.1101_2020.04.19.048751 |
| EPI_ISL_427689 | Centre for Infectious Diseases and Microbiology Public Health | NSW Health Pathology - Institute of Clinical Pathology and Medical Research; Westmead Hospital; University of Sydney | Arnott A et al | Revealing COVID-19 Transmission by SARS-CoV-2 Genome Sequencing and Agent Based Modellin | https://dx.doi.org_10.1101_2020.04.19.048751 |
| EPI_ISL_427760 | Centre for Infectious Diseases and Microbiology Public Health | NSW Health Pathology - Institute of Clinical Pathology and Medical Research; Westmead Hospital; University of Sydney | Sadsad R et al | Revealing COVID-19 Transmission by SARS-CoV-2 Genome Sequencing and Agent Based Modellin | https://dx.doi.org_10.1101_2020.04.19.048751 |
| EPI_ISL_427768 | Centre for Infectious Diseases and Microbiology Public Health | NSW Health Pathology - Institute of Clinical Pathology and Medical Research; Westmead Hospital; University of Sydney | Draper J et al | Revealing COVID-19 Transmission by SARS-CoV-2 Genome Sequencing and Agent Based Modellin | https://dx.doi.org_10.1101_2020.04.19.048751 |
| EPI_ISL_427783 | Centre for Infectious Diseases and Microbiology Public Health | NSW Health Pathology - Institute of Clinical Pathology and Medical Research; Westmead Hospital; University of Sydney | Sim E et al | Revealing COVID-19 Transmission by SARS-CoV-2 Genome Sequencing and Agent Based Modellin | https://dx.doi.org_10.1101_2020.04.19.048751 |
| EPI_ISL_413599 | Centre for Infectious Diseases and Microbiology - Public Health | NSW Health Pathology - Institute of Clinical Pathology and Medical Research; Westmead Hospital; University of Sydney | Timms et al | Revealing COVID-19 Transmission by SARS-CoV-2 Genome Sequencing and Agent Based Modellin | https://dx.doi.org_10.1101_2020.04.19.048751 |
| EPI_ISL_427787 | Centre for Infectious Diseases and Microbiology Public Health | NSW Health Pathology - Institute of Clinical Pathology and Medical Research; Westmead Hospital; University of Sydney | Bachmann N et al | Revealing COVID-19 Transmission by SARS-CoV-2 Genome Sequencing and Agent Based Modellin | https://dx.doi.org_10.1101_2020.04.19.048751 |
| EPI_ISL_427791 | Centre for Infectious Diseases and Microbiology Public Health | NSW Health Pathology - Institute of Clinical Pathology and Medical Research; Westmead Hospital; University of Sydney | Rockett R et al | Revealing COVID-19 Transmission by SARS-CoV-2 Genome Sequencing and Agent Based Modellin | https://dx.doi.org_10.1101_2020.04.19.048751 |
| EPI_ISL_427742 | Centre for Infectious Diseases and Microbiology Public Health | NSW Health Pathology - Institute of Clinical Pathology and Medical Research; Westmead Hospital; University of Sydney | Lam C et al | Revealing COVID-19 Transmission by SARS-CoV-2 Genome Sequencing and Agent Based Modellin | https://dx.doi.org_10.1101_2020.04.19.048751 |
| EPI_ISL_427673 | Centre for Infectious Diseases and Microbiology Public Health | NSW Health Pathology - Institute of Clinical Pathology and Medical Research; Westmead Hospital; University of Sydney | Gray K et al | Revealing COVID-19 Transmission by SARS-CoV-2 Genome Sequencing and Agent Based Modellin | https://dx.doi.org_10.1101_2020.04.19.048751 |
| EPI_ISL_427692 | Centre for Infectious Diseases and Microbiology Public Health | NSW Health Pathology - Institute of Clinical Pathology and Medical Research; Westmead Hospital; University of Sydney | Timms V et al | Revealing COVID-19 Transmission by SARS-CoV-2 Genome Sequencing and Agent Based Modellin | https://dx.doi.org_10.1101_2020.04.19.048751 |
| EPI_ISL_427700 | Centre for Infectious Diseases and Microbiology Public Health | NSW Health Pathology - Institute of Clinical Pathology and Medical Research; Westmead Hospital; University of Sydney | Gall M et al | Revealing COVID-19 Transmission by SARS-CoV-2 Genome Sequencing and Agent Based Modellin | https://dx.doi.org_10.1101_2020.04.19.048751 |
| EPI_ISL_427761 | Centre for Infectious Diseases and Microbiology Public Health | NSW Health Pathology - Institute of Clinical Pathology and Medical Research; Westmead Hospital; University of Sydney | Arnott A et al | Revealing COVID-19 Transmission by SARS-CoV-2 Genome Sequencing and Agent Based Modellin | https://dx.doi.org_10.1101_2020.04.19.048751 |
| EPI_ISL_427795 | Centre for Infectious Diseases and Microbiology Public Health | NSW Health Pathology - Institute of Clinical Pathology and Medical Research; Westmead Hospital; University of Sydney | Sadsad R et al | Revealing COVID-19 Transmission by SARS-CoV-2 Genome Sequencing and Agent Based Modellin | https://dx.doi.org_10.1101_2020.04.19.048751 |
| EPI_ISL_427799 | Centre for Infectious Diseases and Microbiology Public Health | NSW Health Pathology - Institute of Clinical Pathology and Medical Research; Westmead Hospital; University of Sydney | Draper J et al | Revealing COVID-19 Transmission by SARS-CoV-2 Genome Sequencing and Agent Based Modellin | https://dx.doi.org_10.1101_2020.04.19.048751 |
| EPI_ISL_427802 | Centre for Infectious Diseases and Microbiology Public Health | NSW Health Pathology - Institute of Clinical Pathology and Medical Research; Westmead Hospital; University of Sydney | Sim E et al | Revealing COVID-19 Transmission by SARS-CoV-2 Genome Sequencing and Agent Based Modellin | https://dx.doi.org_10.1101_2020.04.19.048751 |
| EPI_ISL_413600 | Centre for Infectious Diseases and Microbiology - Public Health | NSW Health Pathology - Institute of Clinical Pathology and Medical Research; Westmead Hospital; University of Sydney | Gall et al | Revealing COVID-19 Transmission by SARS-CoV-2 Genome Sequencing and Agent Based Modellin | https://dx.doi.org_10.1101_2020.04.19.048751 |
| EPI_ISL_427749 | Centre for Infectious Diseases and Microbiology Public Health | NSW Health Pathology - Institute of Clinical Pathology and Medical Research; Westmead Hospital; University of Sydney | Bachmann N et al | Revealing COVID-19 Transmission by SARS-CoV-2 Genome Sequencing and Agent Based Modellin | https://dx.doi.org_10.1101_2020.04.19.048751 |
| EPI_ISL_427737 | Centre for Infectious Diseases and Microbiology Public Health | NSW Health Pathology - Institute of Clinical Pathology and Medical Research; Westmead Hospital; University of Sydney | Rockett R et al | Revealing COVID-19 Transmission by SARS-CoV-2 Genome Sequencing and Agent Based Modellin | https://dx.doi.org_10.1101_2020.04.19.048751 |
| EPI_ISL_427738 | Centre for Infectious Diseases and Microbiology Public Health | NSW Health Pathology - Institute of Clinical Pathology and Medical Research; Westmead Hospital; University of Sydney | Lam C et al | Revealing COVID-19 Transmission by SARS-CoV-2 Genome Sequencing and Agent Based Modellin | https://dx.doi.org_10.1101_2020.04.19.048751 |
| EPI_ISL_427708 | Centre for Infectious Diseases and Microbiology Public Health | NSW Health Pathology - Institute of Clinical Pathology and Medical Research; Westmead Hospital; University of Sydney | Gray K et al | Revealing COVID-19 Transmission by SARS-CoV-2 Genome Sequencing and Agent Based Modellin | https://dx.doi.org_10.1101_2020.04.19.048751 |
| EPI_ISL_427682 | Centre for Infectious Diseases and Microbiology Public Health | NSW Health Pathology - Institute of Clinical Pathology and Medical Research; Westmead Hospital; University of Sydney | Timms V et al | Revealing COVID-19 Transmission by SARS-CoV-2 Genome Sequencing and Agent Based Modellin | https://dx.doi.org_10.1101_2020.04.19.048751 |
| EPI_ISL_427764 | Centre for Infectious Diseases and Microbiology Public Health | NSW Health Pathology - Institute of Clinical Pathology and Medical Research; Westmead Hospital; University of Sydney | Gall M et al | Revealing COVID-19 Transmission by SARS-CoV-2 Genome Sequencing and Agent Based Modellin | https://dx.doi.org_10.1101_2020.04.19.048751 |
| EPI_ISL_427683 | Centre for Infectious Diseases and Microbiology Public Health | NSW Health Pathology - Institute of Clinical Pathology and Medical Research; Westmead Hospital; University of Sydney | Arnott A et al | Revealing COVID-19 Transmission by SARS-CoV-2 Genome Sequencing and Agent Based Modellin | https://dx.doi.org_10.1101_2020.04.19.048751 |
| EPI_ISL_427775 | Centre for Infectious Diseases and Microbiology Public Health | NSW Health Pathology - Institute of Clinical Pathology and Medical Research; Westmead Hospital; University of Sydney | Sadsad R et al | Revealing COVID-19 Transmission by SARS-CoV-2 Genome Sequencing and Agent Based Modellin | https://dx.doi.org_10.1101_2020.04.19.048751 |
| EPI_ISL_427796 | Centre for Infectious Diseases and Microbiology Public Health | NSW Health Pathology - Institute of Clinical Pathology and Medical Research; Westmead Hospital; University of Sydney | Draper J et al | Revealing COVID-19 Transmission by SARS-CoV-2 Genome Sequencing and Agent Based Modellin | https://dx.doi.org_10.1101_2020.04.19.048751 |
| EPI_ISL_427721 | ACT Pathology, The Canberra Hospital | NSW Health Pathology - Institute of Clinical Pathology and Medical Research; Westmead Hospital; University of Sydney | Sim E et al | Revealing COVID-19 Transmission by SARS-CoV-2 Genome Sequencing and Agent Based Modellin | https://dx.doi.org_10.1101_2020.04.19.048751 |

We gratefully acknowledge all the Authors from the Originating laboratories responsible for obtaining the specimens and the Submitting laboratories where genetic sequence data were generated and shared via the GISAID Initiative, on which this research is based.
Genome sequences and meta-data should be downloaded from https://www.gisaid.org.
